# Mitochondrial dysfunction underlying sporadic inclusion body myositis is ameliorated by the mitochondrial homing drug MA-5

**DOI:** 10.1101/2020.03.17.995159

**Authors:** Yoshitsugu Oikawa, Rumiko Izumi, Masashi Koide, Yoshihiro Hagiwara, Makoto Kanzaki, Naoki Suzuki, Koichi Kikuchi, Tetsuro Matsuhashi, Yukako Akiyama, Mariko Ichijo, Takafumi Toyohara, Takehiro Suzuki, Eikan Mishima, Yasutoshi Akiyama, Yoshiaki Ogata, Chitose Suzuki, Masashi Aoki, Eiji Itoi, Shigeo Kure, Ken-ichiro Hayashi, Takaaki Abe

## Abstract

Sporadic inclusion body myositis (sIBM) is the most common idiopathic inflammatory myopathy, and several reports have suggested that mitochondrial abnormalities are involved in its etiology.

We recruited 9 sIBM patients and found significant histological changes and an elevation of growth differential factor 15 (GDF15), a marker of mitochondrial disease, strongly suggesting the involvement of mitochondrial dysfunction. Bioenergetic analysis of sIBM patient myoblasts revealed impaired mitochondrial function.

Decreased ATP production, reduced mitochondrial size and reduced mitochondrial dynamics were also observed in sIBM myoblasts. Cell vulnerability to oxidative stress also suggested the existence of mitochondrial dysfunction.

Mitochonic acid-5 (MA-5) increased the cellular ATP level, reduced mitochondrial ROS, and provided protection against sIBM myoblast death.

MA-5 also improved the survival of sIBM skin fibroblasts as well as mitochondrial morphology and dynamics in these cells. The reduction in the gene expression levels of Opa1 and Drp1 was also reversed by MA-5, suggesting the modification of the fusion/fission process. These data suggest that MA-5 may provide an alternative therapeutic strategy for treating not only mitochondrial diseases but also sIBM.

## Introduction

Sporadic inclusion body myositis (sIBM) is the most common idiopathic inflammatory myopathy in individuals over the age of 50 years[1], and it is distinguished from other idiopathic inflammatory myopathies by early asymmetric finger flexor and knee extensor weakness, which leads to the loss of hand function and a propensity to fall[2]. To date, its etiology is still unclear, and there is no effective treatment. Recently, there have been several reports showing the involvement of mitochondrial dysfunction in the pathogenesis of sIBM. 1) Endomysial inflammatory infiltrates and rimmed vacuoles are found in muscle biopsy tissues from sIBM patients[3, 4]. Rimmed vacuoles are also commonly detected in muscle tissue from patients with mitochondrial myopathy[5]. 2) Ragged-red fibers and cytochrome oxidase (COX)-negative fibers, both of which are also specific to mitochondrial disease[6], are observed in sIBM[4, 7]. 3) Mitochondrial DNA (mtDNA) deletions have been reported in sIBM[8, 9]. Therefore, these findings suggest the involvement of mitochondrial dysfunction in the etiology of sIBM.

We recently reported a new mitochondria-homing drug, mitochonic acid-5 (MA-5) (4-[2,4-difluorophenyl]-2-[1H-indole-3-yl]-4-oxobutanoic acid), which increases the cellular ATP level and reduces mitochondrial reactive oxygen species (mtROS) production, protecting patients with mitochondrial dysfunction from fibroblast death[10–12]. It also prolongs the survival of a mouse model of mitochondrial disease[11]. This sparked our interest in using MA-5 to treat sIBM. Here, we demonstrated a significant elevation of a mitochondrial disease biomarker, growth differential factor 15 (GDF15)[13], in sIBM patient serum and found mitochondrial dysfunction in both patient myoblasts and skin fibroblasts. Under these conditions, MA-5 improved cell survival, increased ATP, and improved mitochondrial morphology and dynamics, suggesting the potential of MA-5 for sIBM therapy.

## Materials and Methods

### Compounds

MA-5,4-(2,4-difluorophenyl)-2-(1H-indole-3-yl)-4-oxobutanoic acid was chemically synthesized at Okayama University of Science as previously reported[10]. BSO (L-buthionine-[S,R]-sulfoximine) was purchased from Wako Pure Chemical Industries.

### Primary myoblast isolation

The tissue samples were washed with normal saline to remove blood cells and transferred to sterile Dulbecco’s modified Eagle’s medium (DMEM; Wako Pure Chemicals Industries, Osaka, Japan) supplemented with 1% penicillin-streptomycin. The tissue was minced and digested with 0.2% collagenase (Wako Pure Chemicals Industries) and 0.1% DNase I (Sigma-Aldrich, St. Louis, MO) for 1 h at 37 °C. PBS was added to the digested muscle tissue samples, which were filtered through a 70-μm cell strainer (BD Biosciences, Franklin Lakes, NJ) and centrifuged at 700 x g for 20 min. The pellets were resuspended in 1 ml staining solution composed of 1% bovine serum albumin (BSA; Sigma-Aldrich) in PBS and then incubated with an Fc receptor blocking solution (Human TruStain FcX, 1:20 in staining buffer; Biolegend, San Diego, CA) for 10 min. The cells were seeded on 24-well chamber slides coated with Matrigel (Dow Corning, Corning, NY) in growth medium containing DMEM/Ham’s F10 mixture supplemented with 20% fetal bovine serum, 1% penicillin-streptomycin, 1% chicken embryonic extract (United States Biological, Salem, MA), and 2.5 ng/mL basic fibroblast growth factor (ReproCELL, Kanagawa, Japan) and incubated at 37 °C under 5% CO_2_ atmosphere. When the cells reached 80% - 90% confluence, the adherent cells were dissociated by treatment with 1 mM EDTA and passaged into a new Matrigel-coated 10 cm dish[14].

### Patients

Clinical date from five male and four female patients aged 57 to 89 years with a diagnosis of sporadic IBM who were hospitalized between 1995 and 2016 in the Department of Neurology of Tohoku University were evaluated retrospectively. The course of the disease, the distribution of muscle weakness, CK activity, and the results of electrophysiological studies were evaluated. The basis for including the patients was the presence of rimmed vacuoles in muscle biopsy tissue[15].

### Human muscle myoblasts and skin fibroblasts

Myoblasts and fibroblasts obtained from muscle and skin biopsy tissue from sIBM patients were collected at Tohoku University Hospital under the approval of the Ethical Committee of Tohoku University, and written informed consent was obtained from all subjects.

In the cell viability experiments, myoblasts were cultured in 4.5 g/L high-glucose DMEM/Ham’s F-10 with 20% FBS, 1% chick embryo extract (United States Biological), 2.5 ng/mL βFGF (ReproCELL), 1% GlutaMAX (Thermo Fisher Scientific) and 1% sodium pyruvate (Thermo Fisher Scientific) at 37 °C in 5% CO_2_. Fibroblasts were cultured in 1.0 g/L low-glucose DMEM with 10% FBS at 37 °C in 5% CO_2_.

### Immunocytochemical analysis

Myoblasts from sIBM patients were fixed with 4% paraformaldehyde for 15 min at room temperature. Nonspecific binding was blocked by two washes with 1% bovine serum albumin (BSA)/PBS. Myoblasts were then permeabilized with 0.1% Triton X-100/1% BSA/PBS. Then, the myoblasts were incubated overnight at 4 °C with primary antibodies against sarcomeric myosin, MF-20 (Developmental Studies Hybridoma Bank, Iowa City, IA, 1:200) and Desmin (Dako, Santa Clara, CA, 1:100). After washing with PBS, the cells were incubated with biotinylated goat anti-mouse IgG (Sigma, St. Louis, MO, 1:500) as the secondary antibody for an hour. At the end of the incubation time, the cells were washed twice with PBS, and myoblasts were analyzed using Axio Imager 2 microscope (Carl Zeiss, Oberkochen, Germany).

### DNA extraction

Total genomic DNA was extracted from myoblasts from all participating patients using a High Pure PCR Template Preparation Kit (Roche, Basel, Switzerland).

### Mitochondrial DNA amplification

Long-range PCR was performed using PrimeSTAR^®^ GXL DNA Polymerase (R050A) (Takara, Shiga, Japan). A 10,143 kb fragment was amplified by PCR in a thermal cycler (TaKaRa PCR Thermal Cycler) using the following primers: 5’ GAG-GCCTAACCCCTGTCTTT 3’ (forward) and 5’ AGCTTTGGGTGCTAATGGTG 3’ (reverse). The PCR conditions were as follows: initial denaturation at 98 °C for 1 min, 20 cycles of 10 s at 98 °C, 15 s at 68 °C and 10 min at 68 °C and then 25 cycles at 98 °C for 10 s, 56 °C for 15 s, and 68 °C for 10 min. In addition, the purified products were separated by 0.8% acrylamide gel electrophoresis at 100 V for 60 min, and the bands were detected by a BIO-RAD Gel DocTM EZ Imager. The band intensities were analyzed by Image lab software.

### SDS-PAGE and Western blotting

Equal amounts of protein (15 μg) were separated by SDS-PAGE on a 12% gel. The proteins were transferred onto a polyvinylidene difluoride membrane (BIO-RAD, #1704273, Hercules, CA). The membrane was incubated overnight with primary antibodies against total rodent OXPHOS (ab110413, Abcam), MT-NDI (ab181848, Abcam) and β-actin (sc47778, Santa Cruz Biotechnology) and incubated for 1 h with HRP-conjugated secondary antibodies (#32430 and #32460, Invitrogen or Thermo Scientific). The protein bands were detected using the enhanced chemiluminescent plus system.

### Reverse transcription and quantitative PCR

Total RNA was extracted using Sepasol-RNA I Super G (Nacalai Tesque, Kyoto, Japan). cDNA was synthesized using a ReverTra Ace qPCR RT Kit (TOYOBO, Osaka, Japan). Quantitative real-time PCR was performed using a TaqMan Gene Expression Assay (Thermo Fisher Scientific) and SYBR Green PCR reagents according to the manufacturer’s instructions on a StepOnePlus^TM^ Real Time PCR System (Thermo Fisher Scientific). The primers used are listed in Table S2. The cycle threshold (Ct) was calculated using the Ct method. The relative mRNA expression levels were normalized to that of GAPDH.

### Next-generation sequencing

The primers used for amplification of the mitochondrial region are listed in Table S3. The following amplification steps were performed: 98 °C for 1 min followed by 30 cycles at 98 °C for 10 s and 68 °C for 11 min. Fragmentation and library construction of the resulting PCR products were performed using an NEBNext Ultra II FS DNA Library Prep Kit (New England Biolabs, Ipswich, MA) according to the manufacturer’s instructions. The libraries were sequenced on a HiSeq 1500 instrument (Illumina Inc., San Diego, CA), which generated 100-bp paired-end reads. Prior to assembly, adapter sequences were removed from the raw reads, and the trimmed reads were compared with human reference sequences in the GenBank database (GenBank ID: J01415.2; https://www.ncbi.nlm.nih.gov/genbank/). Additionally, the Mitomap database (https://www.mitomap.org/MITOMAP) was used to identify sequence variations of the mitochondrial genome in our samples.

### GDF15 and FGF21

The levels of GDF15 and FGF21 were measured using a Quantikine Human GDF15 ELISA Kit and Quantikine Human FGF21 ELISA Kit (R&D Systems, Minneapolis, MN), respectively.

### Cell viability assay and LDH assay in myoblasts from sIBM patients

In the cell viability assay, myoblasts were plated at a density of 3.0 × 10^3^ cells/150 μl in each well (n=4) of a 96-well flat-bottom culture plate. After 24 h of incubation, BSO and MA-5 or 0.1% DMSO was added, and the cells were cultured for 72 h. Cell viability was measured by Cell Count Regent SF (Nacalai Tesque)[10, 12]. The LDH level was measured by cell counting using an LDH cytotoxicity detection kit (Takara).

### Measurement of mitochondrial function by a flux analyzer

The oxygen consumption rate (OCR) and extracellular acidification rate (ECAR) of myoblasts and fibroblasts from sIBM patients and normal volunteers were measured using XF 24 (Agilent Technologies, Santa Clara, CA). Briefly, myoblasts and fibroblasts were cultured in assay medium (70 mM sucrose, 220 mM mannitol, 10 mM KH_2_PO_4_, 5 mM MgCl_2_, 2 mM HEPES, 1.0 mM EGTA and 0.2% [w/v] fatty acid-free BSA, pH 7.2) without CO_2_ for 60 min. Then, after equilibration, the three respiration rates were measured in myoblasts and fibroblasts following injections of four inhibitors (2 μM oligomycin, 1 μM carbonyl cyanide-p-trifluoromethoxyphenylhydrazone (FCCP), 1 μM rotenone and antimycin) of mitochondrial oxidative phosphorylation (OXPHOS).

### Measurement of ATP production after treatment with DMSO or MA-5

Myoblasts and fibroblasts from sIBM patients were cultured in 96-well plates at a density of 3.0 × 10^3^ cells per well (n=4). The ATP level was measured 3 h after DMSO or MA-5 treatment (10 μM) by an ATP measurement kit (Toyo Ink, Tokyo, Japan).

### Microscopic imaging

To obtain images of mitochondria, cells from sIBM patients were stained with MitoTracker red (a molecular probe, shown in red), and the nuclei were counterstained with 4’,6-diamidino-2-phenylindole (DAPI, shown in blue). The samples were then placed into a confocal microscopy system (Nikon, Tokyo, Japan). Images were acquired and analyzed using NIS-Elements with N-SIM analysis software (Nikon) and ImageJ. Briefly, the MitoTracker red signal in the images was changed to grayscale, and a suitable threshold level that allowed the signal intensity of the mitochondria to be distinguished from the background noise was set in ImageJ software. The lengths of the major and minor axes of the mitochondria were measured using ImageJ. Electron microscopy analysis was performed as previously reported[12, 16]. For live-cell imaging, culture dishes with myoblasts and fibroblasts from sIBM patients stained with MitoTracker Green were placed into a KEYENCE BZ-X700 All-in-one Fluorescence Microscope (KEYENCE, Osaka, Japan). Images were taken for 5 min and converted to movie files using a BZ-X Analyzer (KEYENCE). The movies were analyzed with the video editing analysis software VW-H2MA (KEYENCE) to evaluate cell migration[17].

## Results

### Defects in mitochondrial function in sIBM

We recruited 9 sIBM patients (numbers 1-9), and their clinical characteristics are summarized in Table 1. Patients were clinically diagnosed with sIBM with a negative family history at Tohoku University Hospital (Japan). The mean age of the patients was 74.8 ± 10.2 years, and 55% of the patients were male. The age of onset varied from 47 to 82 years, and the age at diagnosis varied from 54 to 86 years. The most affected muscles were in the shoulder girdle and pelvic girdle. Distal upper limb and distal lower limb muscles were also affected in all patients. The serum creatine kinase (CK) level was moderately elevated to 25 – 720 U/L, up to 20 times above the upper range (laboratory normal value < 34 U/L). Electromyography revealed a myogenic pattern in the biceps brachii and vastus lateralis muscles in 2 patients, a neurogenic pattern in the biceps brachii, vastus lateralis, interosseous, and tibialis anterior muscles in 1 patient, and a myogenic pattern in the tibialis anterior muscle in 1 patient, and a neurogenic pattern in the vastus lateralis muscle in 1 patient. In 4 patients, electromyography was normal. Nerve conduction studies showed no abnormalities, as previously reported[15]. Because histological abnormalities in sIBM patient muscles have been reported[18], we examined muscle samples from all patients (Table 1, representative image in Fig. 1).

**Fig 1.**
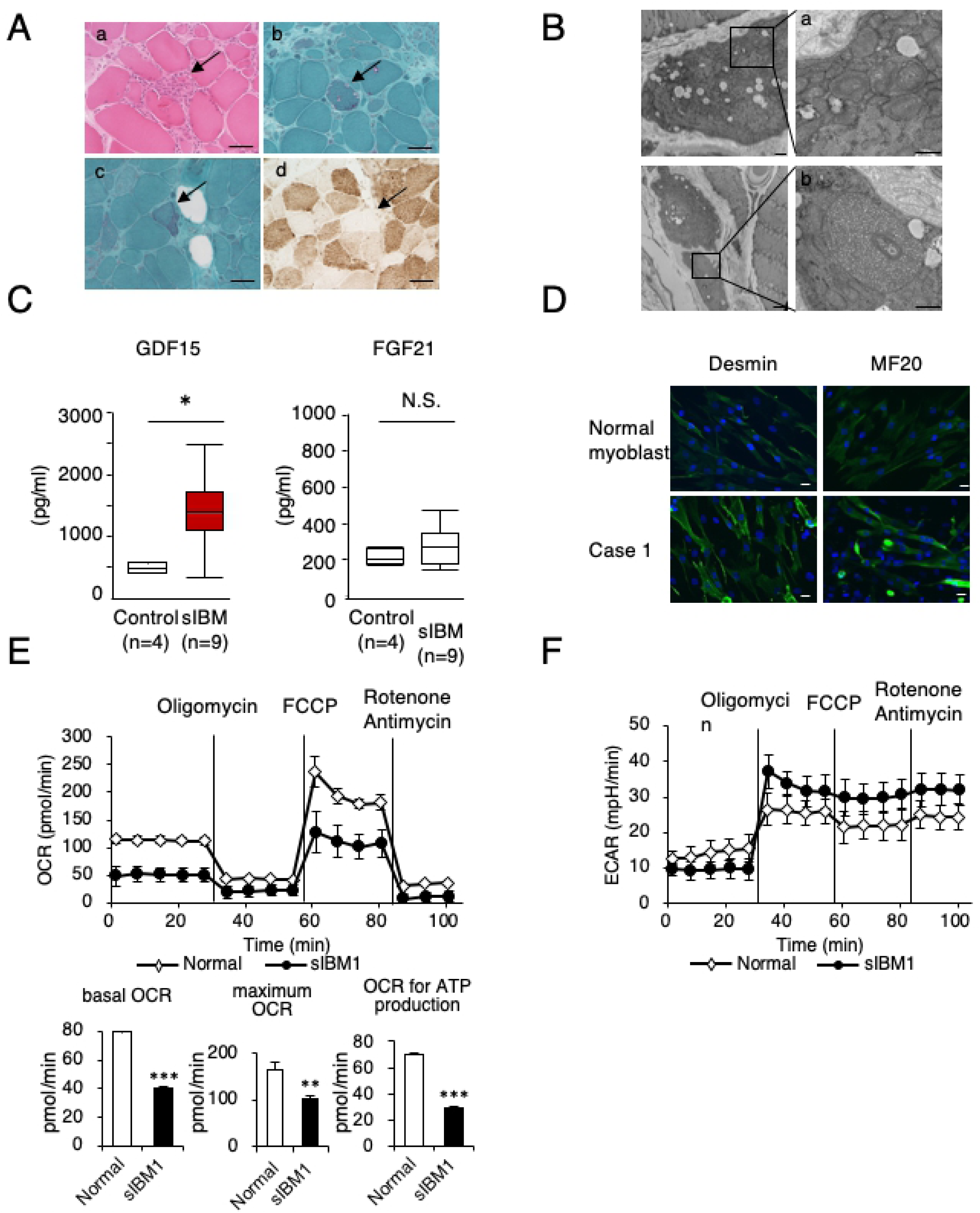
Mitochondrial dysfunction in sIBM patients. **A.** Pathological images of the muscle tissue from sIBM patient (patient 4). **(a)** Hematoxylin-eosin staining of muscle biopsy tissue from a patient with sIBM revealed endomysial inflammatory infiltrates invading nonnecrotic fibers (arrow). **(b)** Modified Gomori trichrome staining of muscle biopsy tissue from a patient with sIBM revealed rimmed vacuoles (arrow). (**c**) Modified Gomori trichrome staining of muscle biopsy tissue from a patient with sIBM revealed ragged-red fibers (arrow). **(d)** The absence of cytochrome c oxidase staining in muscle biopsy tissue from a patient with sIBM (arrow). **B.** Pathological image of muscle tissue from an sIBM patient (patient 4). **(a)** Abnormal mitochondria with concentric cristae were observed by electron microscopy. Scale bar = 5 μm. **(b)** Enlarged mitochondria with abnormal cristae and dense granules. Scale bar = 5 μm. **C.** Measurement of the serum levels of GDF15 and FGF21 in sIBM patients and controls. The data represent the mean ± SEM. \**p*< 0.05 (*p*=0.015, nonparametric Wilcoxon rank sum test; n = 9 patients and n = 4 controls). **D.** Representative immunostaining of sIBM patient tissue and normal myoblasts with desmin and MF20. Scale bars = 50 μm. **E.** Bioenergetic assay of normal and sIBM patient myoblasts (sIBM1) in comparison with the OCR. The data represent the mean ± SD. \*\**p*< 0.01, \*\*\**p*< 0.001 (unpaired two-tailed Student’s *t*-test versus normal myoblasts). **F.** Bioenergetic assay of normal and sIBM patient myoblasts (sIBM1) in comparison with the ECAR. The data represent the mean ± SD.

**Table 1.** Characteristics of sIBM patients. Table shows the characteristics of sIBM patients and the serum levels of GDF15 and FGF21, light microscopy findings and electron microscopy (EM) findings. sIBM, sporadic inclusion body myositis. GDF15, growth differential factor 15. FGF21, fibroblast growth factors 21. RVs, rimmed vacuoles. EIMs, endomysial inflammatory infiltrates invading nonnecrotic fibers. RRFs, ragged red fibers.

| case | Cell ID | sex | Age of onset | Age of diagnosis | Age examined | CK level (U/L) | GDF15 (pg/mL) | FGF21 (pg/mL) | RV | EIM | RRF | COX negative -fiber | Abnormal mitochondria (EM) | Mitochondria DNA large deletion |
| --- | --- | --- | --- | --- | --- | --- | --- | --- | --- | --- | --- | --- | --- | --- |
| 1 | sIBM1 | F | 82 | 86 | 89 | 161 | 2469 | 149 | + |  | + | + | + | none |
| 2 | sIBM2 | F | 75 | 81 | 84 | 417 | 1505 | 382 | + | + | + | + |  | none |
| 3 | sIBM3 | M | 65 | 69 | 71 | 692 | 1175 | 164 | + | + |  |  | + | none |
| 4 |  | M | 60 | 65 | 71 | 619 | 1825 | 275 | + | + | + |  | + |  |
| 5 |  | M | 55 | 61 | 65 | 720 | 327 | 296 | + | + | + | + |  |  |
| 6 |  | M | 64 | 68 | 71 | 419 | 1585 | 327 | + | + | + |  |  |  |
| 7 |  | M | 64 | 68 | 76 | 551 | 1378 | 478 | + | + |  | + |  |  |
| 8 |  | F | 63 | 86 | 89 | 25 | 1025 | 272 | + | + |  | + |  |  |
| 9 |  | F | 57 | 54 | 57 | 242 | 1345 | 204 | + | + |  |  |  |  |

Light microscopy revealed a wide variety of pathological muscle fiber sizes in all patients. Endomysial mononuclear cell infiltration was observed in 8 out of 9 patients, and there were many atrophic or hypertrophic muscle fibers with internal nuclei and splitting; both endomysial inflammation and muscle fiber degeneration were found to be characteristic of sIBM[4] (Table 1). All the samples showed rimmed vacuoles by modified Gomori trichrome (mGT) staining[15] (Table 1, Fig. 1A). Ragged-red fibers[19] were also found in 5 patients (patients 1, 2, 4, 5, and 6; Table 1, Fig. 1A). In addition, deficient cytochrome oxidase (COX) activity[9] was observed in 5 sIBM patients (patients 1, 2, 5, 7, and 8, Table 1, Fig. 1A). By electron microscopy myofibrillar disorganization[15] was observed in 3 patients (patients 1, 3, and 4; Table 1). Two patients (patients 1 and 3; Table 1) had mitochondrial abnormalities with concentric cristae and enlarged mitochondria (Fig. 1B), as previously reported[18]. These data strongly suggest mitochondrial dysfunction in sIBM patient myoblasts.

In the clinical setting, CK[1], amyloidogenic-related molecules (β-secretase 1[20], presenilin-1[21], and soluble Aβ precursor protein[20]) and cytosolic 5’-nucleotidase 1A (cN-1A) autoantibodies[22] are used for the diagnosis of sIBM, but none of these are specific. Recently, GDF15[13, 23] and FGF21[23] have been recognized to be more sensitive and specific diagnostic markers for mitochondrial diseases than the lactate to pyruvate (L/P) ratio. Because our results raise the possibility of the pathophysiological involvement of mitochondrial dysfunction in sIBM, we measured serum GDF15 and FGF21 levels in 9 sIBM patients and 4 normal controls (Table 1, Fig. 1C). We found that the mean serum GDF15 level in sIBM patients was 1404.0 ± 194.0 pg/mL, which was significantly higher than that in normal controls (487.5 ± 45.3 pg/mL; Fig. 1C, left). On the other hand, the serum FGF21 level in sIBM patients was 282.9 ± 35.1 pg/mL, with many sIBM patients having similar values as those of the normal controls (217.0 ± 22.1 pg/mL); there was no significant difference (Fig. 1C; right). These data further suggested underlying mitochondrial dysfunction in sIBM patients and that GDF15 may be a candidate marker for sIBM.

To clarify the mechanism of mitochondrial dysfunction in sIBM, we next examined the mitochondrial bioenergetic function governing the oxygen consumption rate (OCR) and extracellular acidification rate (ECAR) using isolated myoblasts[11, 24, 25]. All isolated cells from sIBM patients (patients 1, 2, and 3) and normal controls were desmin- and MF20-positive and thus possessed the properties of myoblasts[2, 26] (Fig. 1D, S1 Fig). Based on bioenergetic analysis, the basal OCR, maximum OCR and OCR for ATP production were significantly reduced in sIBM myoblasts compared with control myoblasts (representative data from patient 1 in Fig. 1E and from patients 2 and 3 in S2 Fig). At the same time, the ECAR values were significantly increased in sIBM myoblasts, suggesting that activated glycolysis compensated for the impairment of cellular oxidative phosphorylation (OXPHOS) (Fig. 1F).

Because mitochondrial bioenergetic dysfunction results in decreased cellular ATP production, we next measured the cellular ATP levels in sIBM myoblasts. As shown in Fig. 2A, the intracellular ATP level was decreased in sIBM myoblasts compared with normal myoblasts. MA-5 (0.01 – 10 μM) significantly increased the ATP level in sIBM myoblasts (representative data from patient 1 in Fig. 2A and from patients 2 and 3 in S3 Fig). We also examined the bioenergetic changes in sIBM myoblasts induced by MA-5.

**Fig 2.**
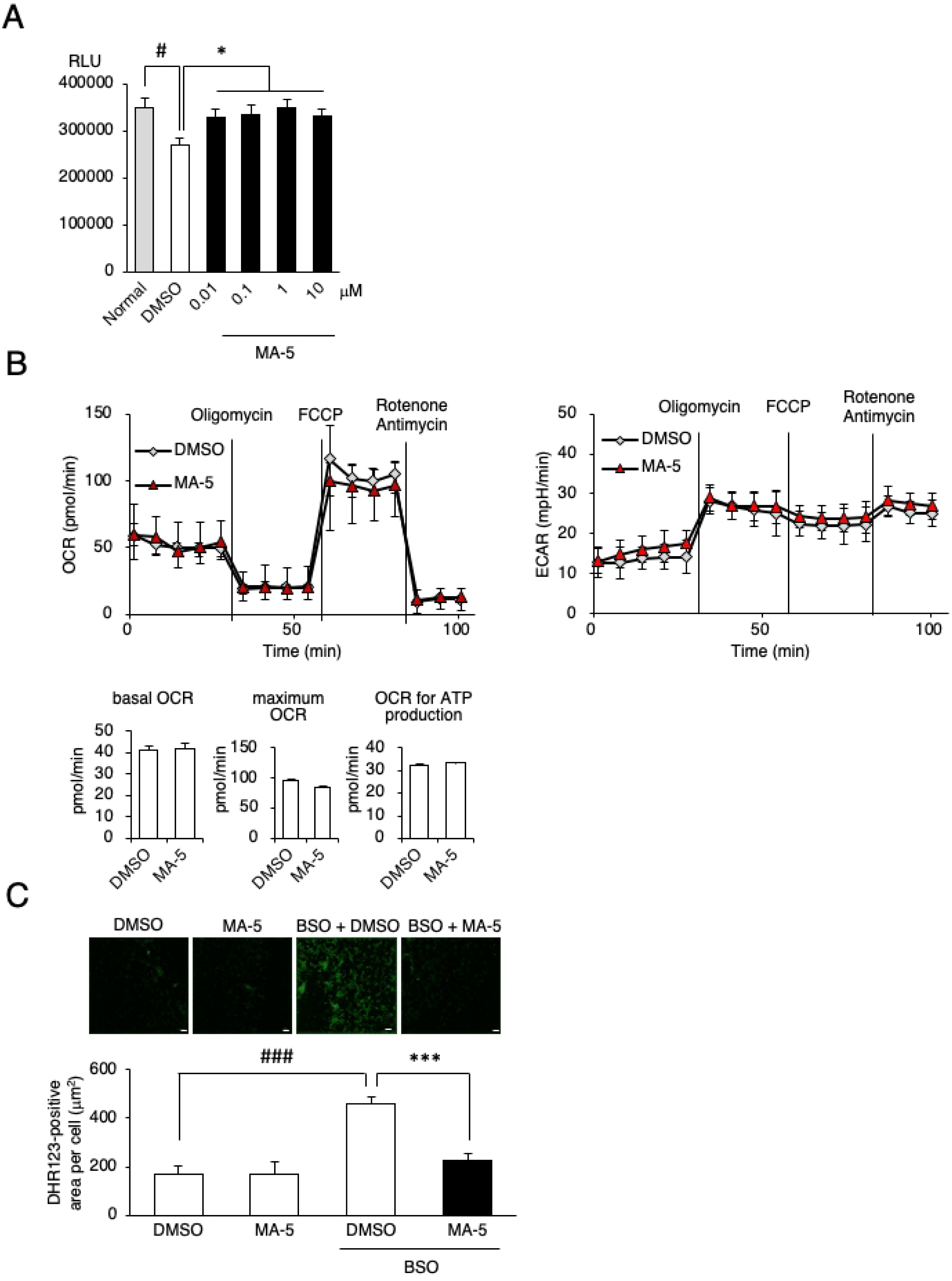
Effect of MA-5 on mitochondrial respiration. **A.** Myoblasts from normal controls and sIBM patients treated with DMSO or MA-5 (0.01, 0.1, 1.0 or 10 μM) for 6 h, and MA-5 increased the ATP level. The data represent the mean ± SE. #*p*< 0.05 (unpaired two-tailed Student’s *t*-test versus normal myoblasts); \**p*< 0.05 (unpaired two-tailed Student’s *t*-test versus DMSO). **B.** Bioenergetic assay of myoblasts from sIBM patients treated with DMSO or MA-5 (10 μM). The data represent the mean ± SD. **C.** MA-5 improved ROS production by BSO. DHR123 staining of sIBM myoblasts treated with DMSO (upper left), MA-5 (upper right), BSO + DMSO (lower left), or BSO + MA-5 (lower right). The data represent the mean ± SE. ###*p*< 0.001 (unpaired two-tailed Student’s *t*-test versus DMSO); \*\*\**p*< 0.001 (unpaired two-tailed Student’s *t*-test versus BSO + DMSO). Scale bars = 200 μm.

Previously, we reported that MA-5 increases ATP without affecting mitochondrial respiration and glycolysis but exerts its function by modifying the inner mitochondrial membrane supercomplex through binding to the inner membrane protein mitofilin[11, 12]. MA-5 increased the cellular ATP (Fig. 2A) without affecting OCR or ECAR values in sIBM myoblasts (Fig. 2B, S4 Fig), which supports our hypothesis of the mechanism of action of MA-5.

To further examine the mechanism of ATP production by MA-5, we measured mitochondrial ROS (mtROS). Mitochondria are the primary source of ROS[27], and if MA-5 induces the overwork of damaged mitochondria to produce ATP, mtROS would be increased. It is well known that mtROS is induced by the glutathione synthesis inhibitor L-buthionine-(S, R)-sulfoximine (BSO)[28], so we added BSO to the culture medium[12]. The mtROS level in sIBM myoblasts was significantly increased by BSO compared with that in normal myoblasts (representative data from patient 1 in Fig. 2C and from patients 2 and 3 in S5 Fig). MA-5 ameliorated the mtROS level (Fig. 2C). These data suggest that MA-5 increased cellular ATP without inducing mitochondrial overwork caused by impaired OXPHOS.

### MA-5 improved the morphology and dynamics of damaged mitochondria

It is well known that morphological changes promote mitochondrial fragmentation, disturb mitochondrial dynamics, and lower the OXPHOS level[29, 30], which likely impairs their capacity to respond to stress. Indeed, fibroblasts of mitochondrial disease patients show mitochondrial dysfunction with network fragmentation[31] and impaired dynamics[32], even under baseline conditions. Thus, these abnormalities are enhanced by oxidative stress and result in the vulnerability of patient fibroblasts.

Therefore, we examined mitochondrial morphology and dynamics in sIBM myoblasts. The mitochondrial network of sIBM myoblasts was broken into smaller pieces and showed a granular pattern, resulting in a reduced mitochondrial area (representative data from patient 1 in Fig. 3A, left), upon treatment with BSO. MA-5 significantly restored the damaged area and reticular morphology of the mitochondria (Fig. 3A, right). Based on electron microscopy, the mitochondrial cristae in sIBM myoblasts were shortened and enlarged to the extent that the cristae junctions were broken, resulting in a reduced mitochondrial length/width ratio (representative data from patient 1 in Fig. 3B). MA-5 also increased the length/width ratio compared to that of the control (Fig. 3B right, S6 Fig).

**Fig 3.**
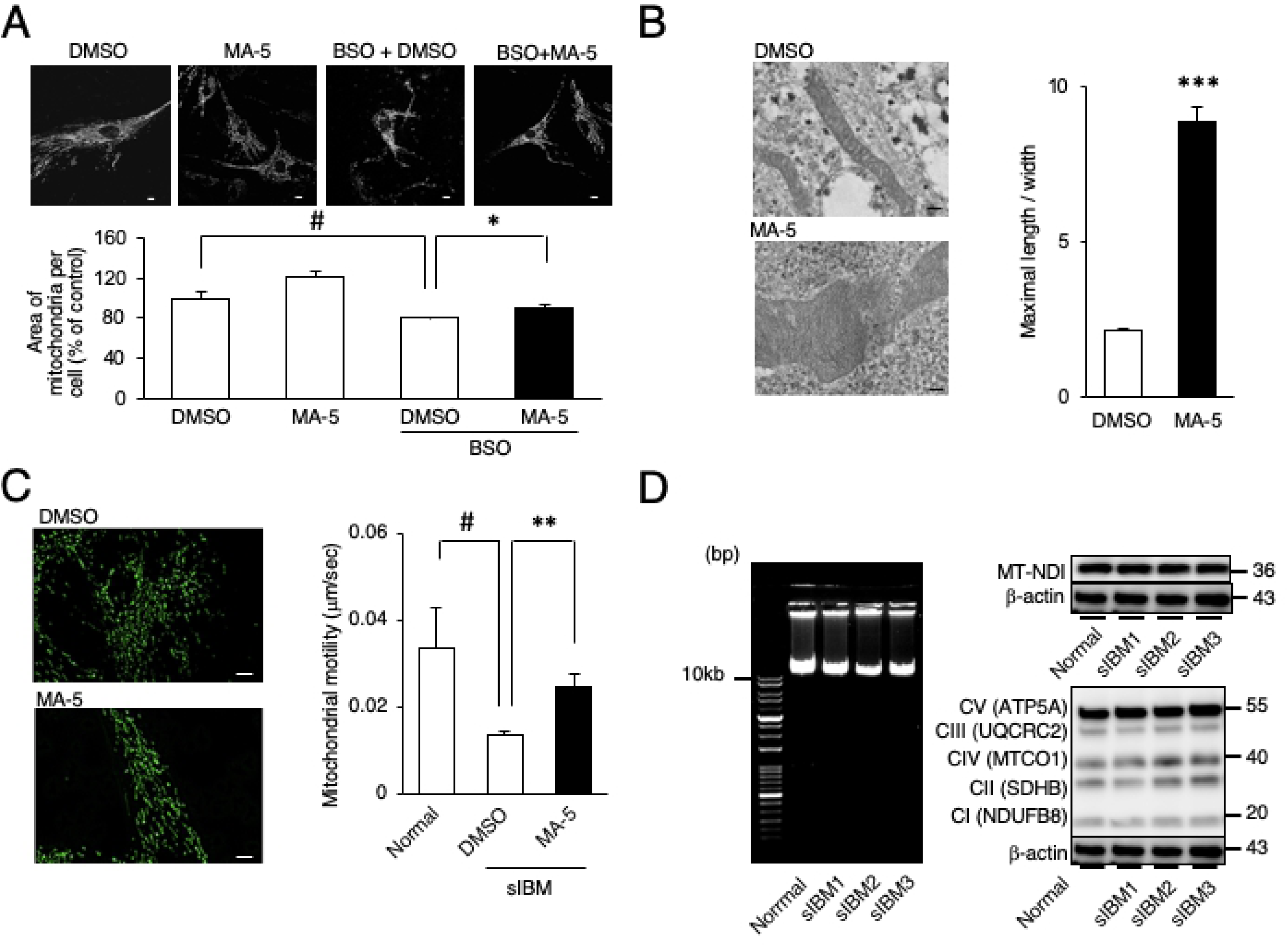
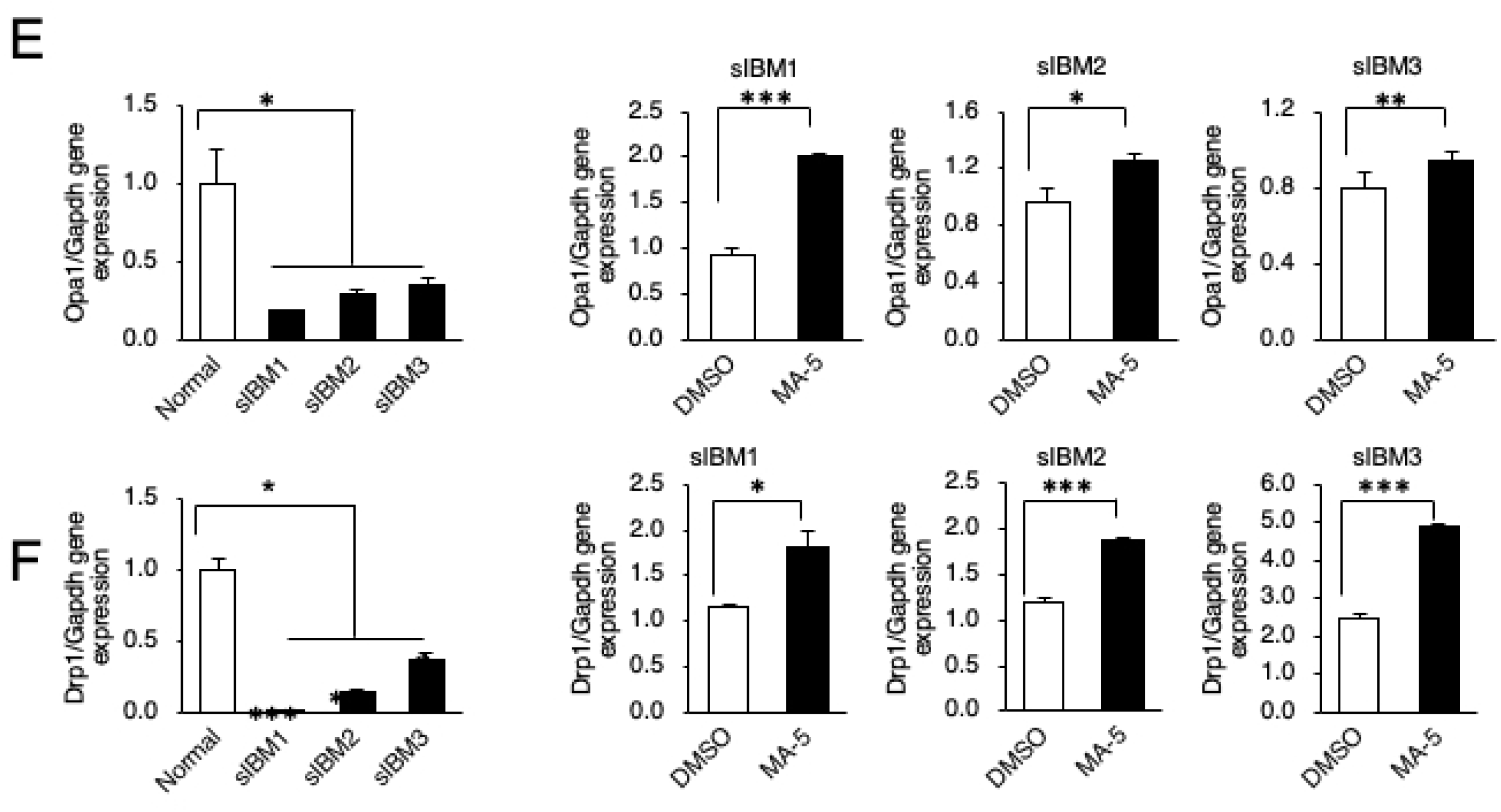
Effect of MA-5 on sIBM myoblasts. **A.** MA-5 improved mitochondrial fragmentation. Structural changes in mitochondria were observed in DMSO, MA-5, BSO + DMSO, BSO + MA-5 myoblasts (300 μM BSO, 0.1% DMSO and 10 μM MA-5) using confocal microscopy imaging. The data represent the mean ± SE. ###*p*< 0.001 (unpaired two-tailed Student’s *t*-test versus DMSO); \*\*\**p*< 0.001 (unpaired two-tailed Student’s *t*-test versus BSO + DMSO). Scale bars = 10 μm. **B.** MA-5 improved cristae length. Note that cristae were damaged and that MA-5 lengthened and tightened the cristae. The ratio of cristae length to width was calculated (n = 100). The data represent the mean ± SE. \*\*\**p*< 0.001 (two-way ANOVA test versus DMSO). Scale bars = 200 nm**. C.** MA-5 improved mitochondrial motility. Mitochondrial motility in sIBM myoblasts treated with DMSO (upper) or MA-5 (lower) was measured. Note that mitochondrial motility was significantly increased by MA-5. The data represent the mean ± SE. #*p*< 0.05 (unpaired two-tailed Student’s *t*-test versus normal myoblasts); \*\**p*< 0.01 (unpaired two-tailed Student’s *t*-test versus sIBM myoblasts treated with DMSO). Scale bars = 10 μm**. D.** Long-range PCR analysis of mtDNA from individual myoblasts from normal controls and sIBM patients (left). Western blotting of individual myoblasts from normal controls and sIBM patients (right) for mitochondrial ND-1 and OXPHOS complexes (CI – CV). β-actin was used as a loading control. **E.** Opa1 was assessed at the transcript level in myoblasts from normal controls and sIBM patients. Significant differences were found in Opa1 mRNA levels in sIBM myoblasts compared with normal control myoblasts. The data represent the mean ± SE. \**p*< 0.05 (two-way ANOVA and Tukey-Kramer test versus normal myoblasts). MA-5 significantly increased Opa1 expression in all patients. \**p*< 0.05, \*\**p*< 0.01, \*\*\**p*< 0.001 (two-way ANOVA and Tukey-Kramer test versus DMSO). **F.** Drp1 was assessed at the transcript level in myoblasts from normal controls and sIBM patients. Significant differences were found in Drp1 mRNA levels in sIBM myoblasts compared with normal control myoblasts. The data represent the mean ± SE. \**p*< 0.05, \*\**p*< 0.01 \*\*\**p*< 0.001 (two-way ANOVA and Tukey-Kramer test versus normal myoblasts). MA-5 significantly increased Drp1 expression in all patients. \**p*< 0.05, \*\*\**p*< 0.001 (two-way ANOVA and Tukey-Kramer test versus DMSO).

We also evaluated mitochondrial dynamics in sIBM myoblasts through time-lapse imaging (Fig. 3C). Mitochondria in sIBM myoblasts were slower (0.0137 ± 0.0011 μm/s) than those in normal myoblasts (0.0336 ± 0.0093 μm/s) (representative data from patient 1 in Fig. 3C, left). MA-5 significantly improved mitochondrial mobility in sIBM myoblasts (0.0279 ± 0.0015 μm/s; Fig. 3C, right; live imaging is shown in S1 Video Data). These data suggest that MA-5 changes the shape of cristae after improving mitochondrial morphology and dynamics in sIBM myoblasts.

Recently, it was reported that mtDNA deletion[33] and mitochondrial DNA rearrangements occur in sIBM[8]. To further identify the molecular mechanism of sIBM, we amplified whole mitochondrial DNA (mtDNA) from myoblasts from normal controls and 3 sIBM patients (patients 1, 2, and 3). Based on PCR amplification of whole mtDNA, no major deletion was identified (band size of ∼10 kb; Fig. 4D, left). We next performed whole mtDNA sequencing by next-generation sequencing (NGS). We found three SNPs: m.1438A>G or A>C in all three patients, m.1382A>C in one patient (patient 1) and m3849G>A in one patient (patient 2)[34–36] (S1 Table). Based on the database search, all mutations were recognized as nonpathogenic (S1 Table).

**Fig. 4.**
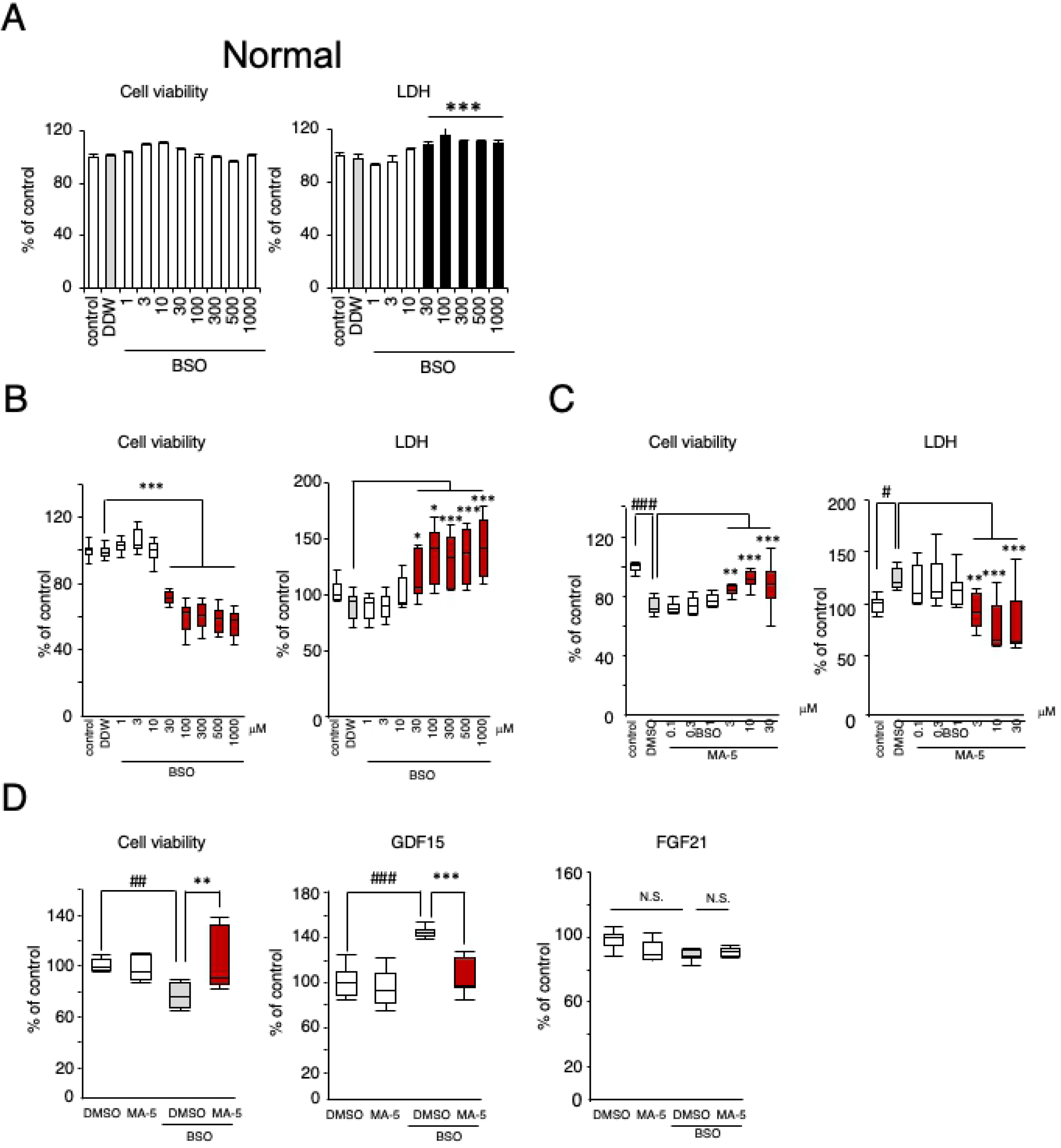
Cell protective effect of MA-5 on sIBM myoblasts. **A.** Sensitivity of normal control myoblasts to BSO. The data represent the mean ± SE. \*\*\**p*< 0.001 (two-way ANOVA and Tukey-Kramer test versus DDW). **B.** Sensitivity of sIBM patient myoblasts to BSO, as measured by the cell viability assay and the level of LDH in the culture medium. The data represent the mean ± SEM. \**p*< 0.05, \*\*\**p*< 0.001 (two-way ANOVA and Tukey-Kramer test versus BSO + DDW). The red square indicates a significant increase or decreased compared with DDW. **C.** Cell protective effect of MA-5 on sIBM myoblasts, as measured by the cell viability assay and the level of LDH in the culture medium. The data represent the mean ± SEM. #*p*< 0.05, ###*p*< 0.001 (two-way ANOVA and Tukey post hoc test versus control); \*\**p*< 0.01, \*\*\**p*< 0.001 (two-way ANOVA and Tukey-Kramer test versus BSO + DMSO). The red square indicates a significant increase or decrease compared with DMSO. **D.** Measurement of sIBM patient myoblasts by the cell viability assay (left panel) after 72-h DMSO application as a control or 72-h 10 μM MA-5 treatment under oxidative stress conditions induced by 24-h BSO treatment. The levels of GDF15 (middle panel) in the medium of cultured myoblasts were measured in the same manner. The levels of FGF21 (right panel) in the medium of cultured myoblasts were measured in the same manner. The data represent the mean ± SE. ##*p*< 0.01, ###*p*< 0.001 (two-way ANOVA and Tukey-Kramer test versus DMSO); ** *p*< 0.01, \*\*\**p*< 0.001 (two-way ANOVA and Tukey-Kramer test versus BSO + DMSO).

However, among these, m.1438A>G and A>C have been reported to be associated with type 2 diabetes[37], schizophrenia[38], cystic fibrosis and aminoglycoside ototoxicity in patients[34], all of which have been reported to have a relationship with mitochondrial dysfunction.

Because m.1438 is located upstream of NADH dehydrogenase 1 (ND1), we examined the protein expression level of ND1 in sIBM myoblasts. As shown in Fig. 3D (right), we could not find any difference in the ND1 protein expression level. We also examined the expression level of respiratory complex I – V proteins: complex I (NDUFB8), complex II (SDHB), complex III (UQCRC2), complex IV (MTCO1) and complex V (ATP5A). However, we could not find any of the significant differences (Fig. 3D, left).

Mitochondrial fission and fusion regulate a number of cellular processes, including bioenergetic potency, mitochondrial apoptosis, autophagy, organelle distribution and morphogenetic processes[39]. Mitochondrial fusion/fission is primarily regulated by dynamin-like GTPase, optic atrophy 1 (Opa1) and dynamin-related protein 1 (Drp1, mitofusin 1/2 (Mfn1/2)[40–42]. Recently, it was reported that the mRNA expression level of Opa1 is reduced in sIBM[43]. It was also reported that OPA1 functionally interacts with MIC60/mitofilin [44]. Based on these findings, we next examined the expression levels of Opa1 and Drp1 in sIBM myoblasts. The Opa1 mRNA level was significantly decreased in sIBM patient myoblasts compared with normal control myoblasts (Fig. 3E). MA-5 significantly increased the Opa1 expression level in all patient myoblasts (Fig. 3E). On the other hand, the expression level of mitofusin 2 (mfn2) was not changed (S7 Fig). We also examined the expression level of Drp1. The Drp1 mRNA level was also decreased in sIBM patient myoblasts compared with normal control myoblasts, and MA-5 significantly increased the reduced expression of Drp1 (Fig. 3F). These data strongly suggest underlying dysfunction in the mitochondrial fusion/fission process in sIBM myoblasts and that MA-5 may improve the damage and/or imbalance.

### MA-5 protected against myoblast cell death

It is well known that compared with those from normal controls, fibroblasts from mitochondrial patients are vulnerable to oxidative stress, such as that induced by BSO[10–12, 28]. As previously suggested, in normal myoblasts, neither cell viability nor the LDH level in the culture medium was affected by BSO (1 – 1000 μM) (Fig. 4A). In contrast, in sIBM myoblasts, cell viability (patients 1, 2, and 3) was significantly reduced by BSO in a dose-dependent manner (combined data is shown in Fig. 4B, left, and patient data is shown in S8 Fig). The mean IC50 value was 1014.8± 413.5 μM, which is in the same range as that in mitochondrial disease fibroblasts (Leigh, MELAS and Kearns-Sayre syndromes), as we previously reported[10, 12]. The LDH level in the culture medium of sIBM patient myoblasts was also increased by BSO (Fig. 4B, right and S8 Fig). The BSO-induced cell death in sIBM myoblasts was significantly ameliorated by MA-5 in a dose-dependent manner (3 – 30 μM in Fig. 4C, left and Fig. S9). The elevated LDH level was also reduced by MA-5 (Fig. 4C, right and S9 Fig). These data support the presence of mitochondrial dysfunction in sIBM myoblasts and the effectiveness of MA-5.

To evaluate whether GDF15 is an effective marker, we examined GDF15 and FGF21 levels in the culture medium of sIBM myoblasts (Fig. 4D). The GDF15 level was significantly increased by BSO, and cell viability was decreased (Fig. 4D, left). MA-5 significantly ameliorated the BSO-induced GDF15 level and increased cell viability in sIBM myoblasts (Fig. 4D, middle). On the other hand, FGF21 was not changed by BSO with or without MA-5 (Fig. 4D, right). These data suggest that the GDF15 level is a useful marker that can predict mitochondrial damage in sIBM patients and evaluate the efficacy of MA-5 treatment.

### Skin fibroblasts as an alternative material for analyzing mitochondrial function in sIBM

One of the difficulties in the diagnosis and analysis of sIBM is the complicated method required to isolate and purify myoblasts from patient muscle specimens[14]. To overcome this, we simultaneously collected skin fibroblasts from sIBM patients and examined them.. To measure mitochondrial bioenergetic function in sIBM patient fibroblasts, we evaluated the basal OCR and maximum OCR values and found that they were reduced (Fig. 5A). The ECAR values were increased in fibroblasts from sIBM patients. MA-5 did not affect the OCR or ECAR values as it did in myoblasts (Fig. 5B).

**Fig 5.**
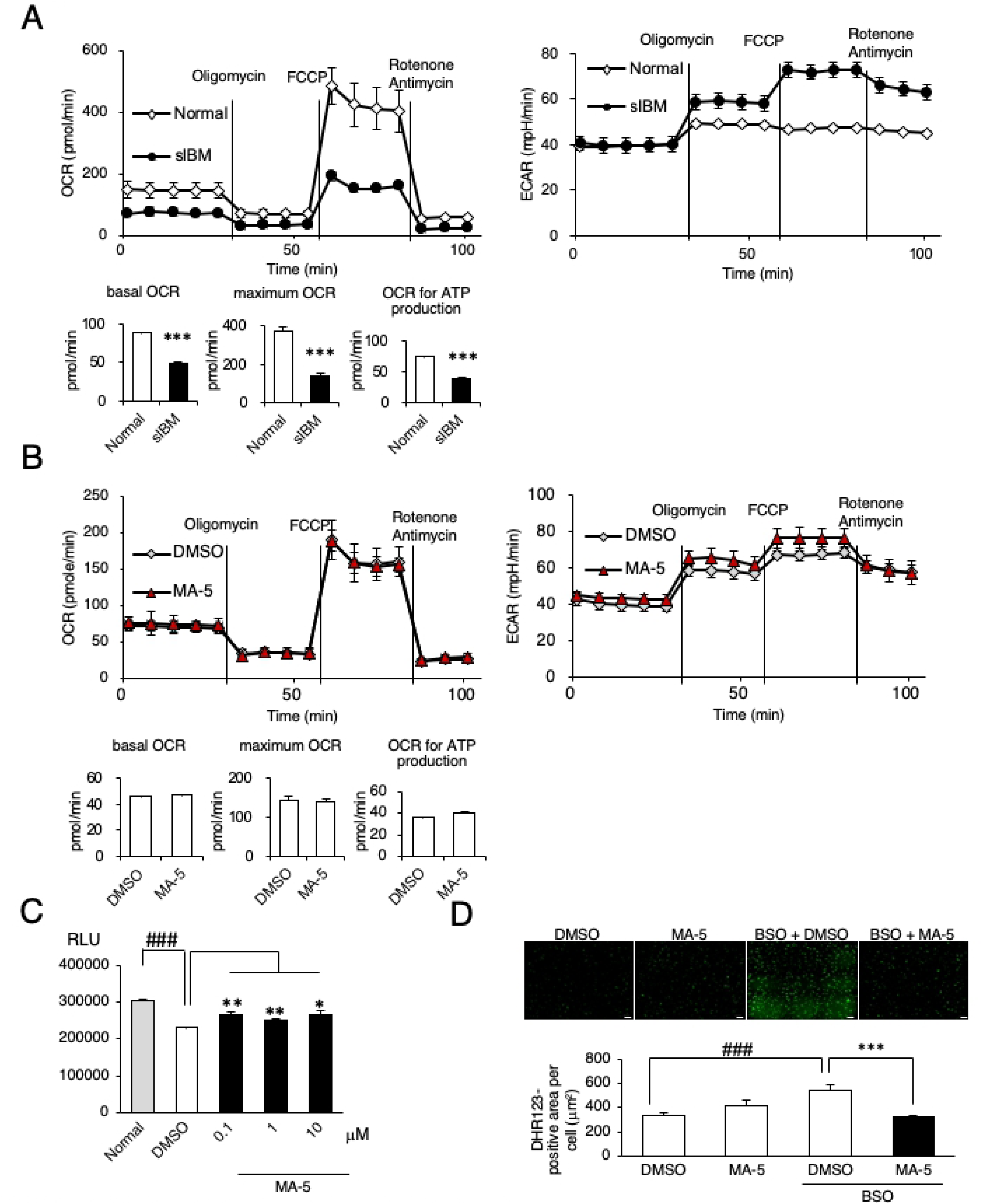
Effect of MA-5 on mitochondrial respiration in sIBM skin fibroblasts. **A.** MA-5 increased the ATP level of fibroblasts from normal controls and sIBM patients treated with DMSO or MA-5 (0.1, 1.0 or 10 μM) for 6 h. The data represent the mean ± SE. ###*p*< 0.001 (unpaired two-tailed Student’s *t*-test versus normal fibroblasts); \**p*< 0.05, \*\**p*< 0.01 (unpaired two-tailed Student’s *t*-test versus DMSO). **B.** MA-5 improved ROS production by BSO. DHR123 staining of sIBM fibroblasts treated with DMSO, MA-5, BSO + DMSO, or BSO + MA-5. The data represent the mean ± SE. ###*p*< 0.001 (unpaired two-tailed Student’s *t*-test versus DMSO); \*\*\**p*< 0.001 (unpaired two-tailed Student’s *t*-test versus BSO + DMSO). Scale bars = 200 μm. **C.** Bioenergetic assay of normal and sIBM patient fibroblasts (sIBM1) in comparison with the OCR (left) and ECAR (right). The data represent the mean ± SD. **D.** Bioenergetic assay of fibroblasts from sIBM patients treated with DMSO or MA-5 (10 μM). The data represent the mean ± SD.

MA-5 (0.1 – 10 μM) significantly increased the ATP level in sIBM patient fibroblasts (representative data from patient 1 in Fig. 5C). Mitochondrial ROS were increased by BSO, and the upregulation of mitochondrial ROS was significantly ameliorated by MA-5 (Fig. 5D). In addition, impaired reticular morphology of the mitochondrial network was also observed in sIBM fibroblasts, and MA-5 improved this morphology compared with that in the BSO group (Fig. 6A). Mitochondrial cristae in the patient fibroblasts were shortened and loosened to the extent that the cristae junctions were broken (Fig. 6B), and MA-5 significantly increased the length/width ratio compared to that in the control (Fig. 6B, S10 Fig). Furthermore, mitochondrial dynamics in sIBM fibroblasts was reduced (0.0101 ± 0.0015 μm/s) compared to that in normal fibroblasts (0.0408 ± 0.0034 μm/s, Fig. 6C), and MA-5 significantly improved mitochondrial mobility in sIBM fibroblasts (0.0253 ± 0.0032 μm/s; live imaging is shown in S2 Video Data). These data were similar to those from sIBM patient myoblasts, suggesting that skin fibroblasts can serve as a comparable tool for diagnostic use in sIBM patients.

**Fig 6.**
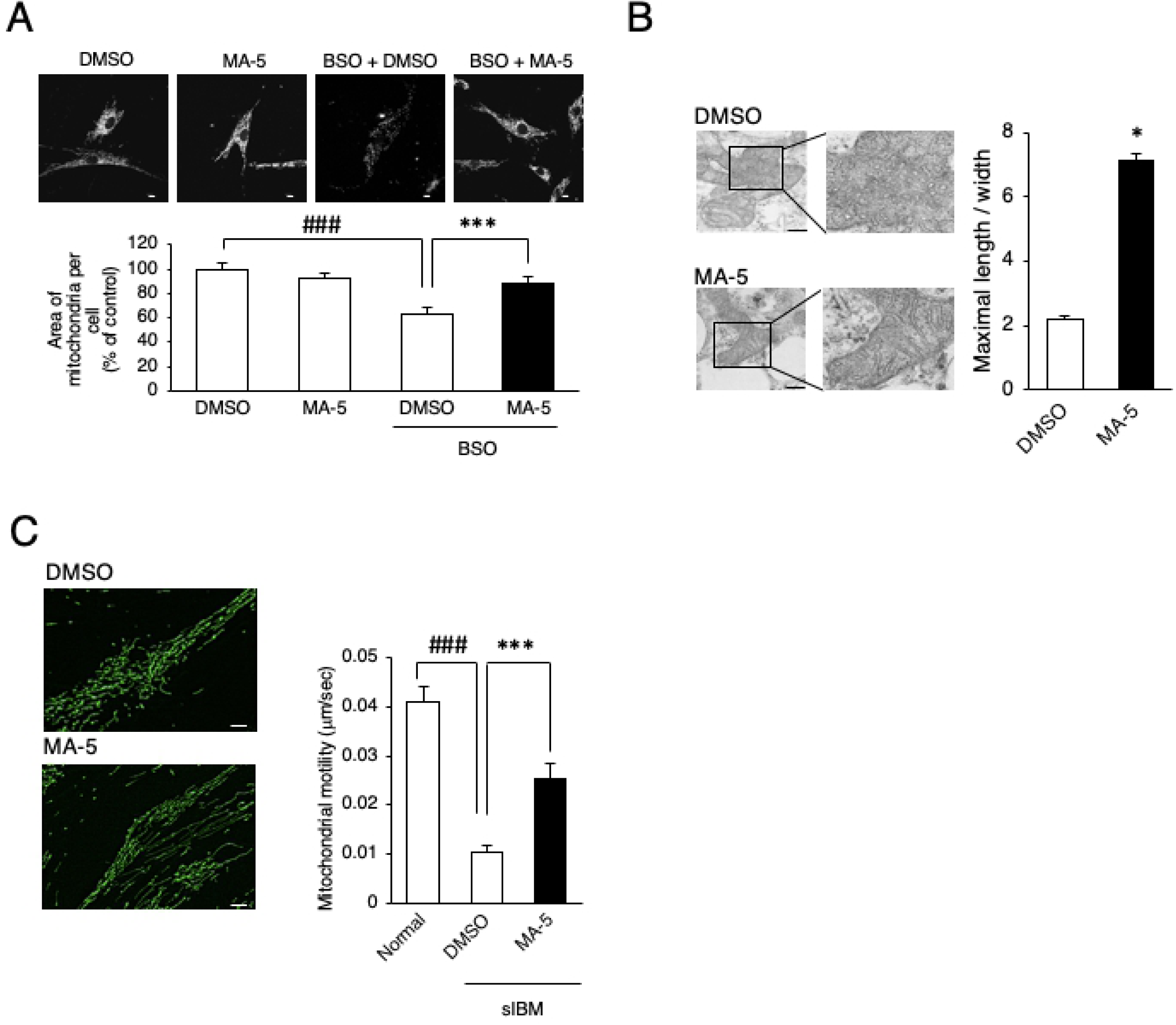
Effect of MA-5 on mitochondrial dynamics in sIBM skin fibroblasts. **A.** MA-5 improved mitochondrial fragmentation. Structural changes in mitochondria were observed in DMSO, MA-5, BSO + DMSO, BSO + MA-5 fibroblasts (300 μM BSO, 0.1% DMSO and 10 μM MA-5) using confocal microscopy imaging. The data represent the mean ± SE. ###*p*< 0.001 (unpaired two-tailed Student’s *t*-test versus DMSO); \*\*\**p*< 0.001 (unpaired two-tailed Student’s *t*-test versus BSO + DMSO). Scale bars = 10 μm. **B.** MA-5 improved cristae length. Note that the cristae were damaged and that MA-5 lengthened and tightened the cristae. The ratio of cristae length to width was calculated (n = 100). The data represent the mean ± SE. \**p*< 0.05 (two-way ANOVA test versus DMSO). Scale bars = 200 nm**. C.** MA-5 improved mitochondrial motility. Mitochondrial motility in sIBM myoblasts treated with DMSO (upper) or with MA-5 (lower) was measured. Note that mitochondrial motility was significantly increased by MA-5. The data represent the mean ± SE. ###*p*< 0.001 (unpaired two-tailed Student’s *t*-test versus normal fibroblasts); \*\*\**p*< 0.001 (unpaired two-tailed Student’s *t*-test versus sIBM fibroblasts treated with DMSO).

## Discussion

Here, we report the presence of mitochondrial dysfunction in sIBM and that a mitochondria-homing drug, MA-5, may be a potential candidate drug for sIBM. Although the abnormal accumulation of aggregated protein within skeletal muscle fibers is considered the pathological cause of sIBM[18], mitochondrial abnormalities are also recognized in sIBM patients.

Previously, an anti-cytosolic 5’-nucleotidase 1A (cN-1A) antibody was found in sIBM patient serum, but the antibody is not highly sensitive[22, 45]. We found that the circulating GDF15 level was significantly higher in sIBM patients than that in normal controls. We also found that the GDF15 level in myoblast culture medium was increased by BSO and decreased by MA-5. These data strongly suggest underlying mitochondrial dysfunction in sIBM patients and that GDF15 may serve as an alternative marker for diagnosing sIBM and evaluating the efficacy of therapies.

We also confirmed bioenergetic mitochondrial dysfunction in sIBM myoblasts. In sIBM patient myoblasts compared with normal myoblasts, we found decreased OCR (Fig. 1E) and reduced ATP production (Fig. 2A). These defects in mitochondrial function result in deficient COX activity[46] and an increased frequency of ragged-red fibers[18, 47] in sIBM myoblasts. This impairment of mitochondrial bioenergetics function could further induce the vulnerability of sIBM myoblasts to oxidative stress (Fig. 4).

Recently, we reported that a synthetic indole compound, MA-5, increased ATP levels via a unique mechanism[10–12]. MA-5 binds to an inner mitochondrial membrane protein, mitofilin, and increases ATP by inducing a conformational change in cristae without changing the membrane potential[12]. We also showed that MA-5 rescues BSO-mediated cellular damage in many types of mitochondrial diseases (MELAS syndrome, Leigh syndrome, KSS, DOA, etc.)[12]. We found that reductions in mitochondrial size (Fig. 3A), the mitochondrial network (Fig. 3B) and mitochondrial dynamics (Fig. 3C) were found in sIBM myoblasts and that MA-5 restored these deficiencies. We also found that the expression levels of both Opa1 (Fig. 3E) and Drp1 (Fig. 3F) were reduced in sIBM myoblasts and that the reductions in these levels were reversed by MA-5 in sIBM myoblasts. Mitochondrial dynamics are important for mitochondrial function, such as organelle recycling and turnover, and an imbalance of the proper ratios of fusion and fission events has recently been found to be associated with many diseases[48, 49]. Opa1 mutations are associated with reduced ATP, and the loss of Opa1 also leads to mitochondrial fragmentation[50] and reduces the percentage of mobile mitochondria[42]. Mutations in Drp1 are associated with a range of neurogenerative disorders[51], and reduced expression levels of Drp1 have been found in cardiac and skeletal muscles during aging[52]. The muscle-specific loss of Drp1 also induces muscle wasting and weakness[53]. Interestingly, both fusion and fission machinery are suppressed in aging sarcopenia, cachexia, and chemotherapy-induced muscle wasting, and Opa1 and Drp1 double-knockout mice show muscle loss[54].

Therefore, MA-5 may in part ameliorate the imbalance in the mitochondrial fusion/fission process and dynamics associated with mitochondrial dysfunction. MA-5 is a novel treatment not only for sIBM but also for sarcopenia, cachexia and muscle atrophy. However, there are some reports that the overexpression of Opa1 ameliorates the phenotype of mitochondrial disease models[39] while the inhibition of Drp1-mediated mitochondrial fission improves mitochondrial dynamics and bioenergetics, stimulating neurogenesis[55]. Therefore, further experiments are necessary to clarify the role of MA-5 in fission/fusion dynamics.

In conclusion, MA-5 is an alternative therapeutic strategy for treating mitochondrial diseases as well as sIBM. The use of GDF15 for diagnostics will also be useful in a forthcoming clinical trial of MA-5.

## Acknowledgement

We thank Mr. Brent Bell and Springer Editing Serive for English editing.

## Author contributions

Yoshitsugu Oikawa and Takaaki Abe designed the experiment. Rumiko Izumi observed muscular tissues by light microscopy and electron microscopy. Masashi Koide and Yoshihiro Hagiwara isolated and purified myoblasts from sIBM patients. Makoto Kanzaki, Tetsuro Matsuhashi, Koichi Kikuchi, Yukako Akiyama, Mariko Ichijo, Takafumi Toyohara, Takehiro Suzuki, Eikan Mishima, Yasutoshi Akiyama, Yoshiaki Ogata, and Chitose Suzuki performed the cell culture experiments. Naoki Suzuki supplied the patient specimens. Masashi Aoki, Eiji Itoi, Shigeo Kure^1^, Ken-ichiro Hayashi and Takaaki Abe discussed the experiment.

## Abbreviations

MA-5: 4-(2,4-difluorophenyl)-2-(1H-indole-3-yl)-4-oxobutanoic acid

MELAS: myopathy encephalopathy

sIBM: sporadic inclusion body myositis

GDF15: growth differential factor 15

COX: cytochrome oxidase

mtDNA: mitochondrial DNA

mtROS: mitochondrial reactive oxygen species

CK: creatine kinase

mGT: Gomori trichrome

OCR: oxygen consumption rate

ECAR: extracellular acidification rate

OXPHOS: oxidative phosphorylation

BSO: L-buthionine-(S, R)-sulfoximine

ND1: NADH dehydrogenase 1

Opa1: optic atrophy 1

Drp1: dynamin-related protein 1

## Supplementary Video Legends

**Video S1** Time-lapse imaging of mitochondria in sIBM myoblasts. sIBM myoblasts were treated with 0.1% DMSO (upper) or 10 μM MA-5 (lower) for 24 h.

**Video S2** Time-lapse imaging of mitochondria in sIBM fibroblasts. sIBM fibroblasts were treated with 0.1% DMSO (upper) or 10 μM MA-5 (lower) for 24 h.

## Supporting Information

**Figure S1**

Immunostaining of sIBM patient myoblasts. (a) Normal myoblasts. (b) sIBM1 myoblasts. (c) sIBM2 myoblasts. (d) sIBM3 myoblasts. Scale bars = 50 μm.

**Figure S2**

Mitochondrial respiration in sIBM myoblasts. Bioenergetic assay of normal and sIBM patient (sIBM2 and sIBM3) myoblasts in comparison with the OCR (left) and ECAR (right). The data represent the mean ± SEM. \**p*< 0.05, \*\*\**p*< 0.001 (unpaired two-tailed Student’s *t*-test versus normal myoblasts).

**Figure S3**

Effect of MA-5 on cellular ATP. Myoblasts from normal controls and sIBM patients (patient 2 and patient 3) were treated with DMSO and MA-5 (0.01, 0.1, 1.0 or 10 μM) for 6 h, and MA-5 increased the ATP level. The data represent the mean ± SEM. #*p* < 0.05, ##*p*< 0.01 (unpaired two-tailed Student’s *t*-test versus normal myoblasts); \**p* < 0.05, \*\**p*< 0.01 (unpaired two-tailed Student’s *t*-test versus DMSO).

**Figure S4**

Effect of MA-5 on mitochondrial respiration. Bioenergetic assay of myoblasts from sIBM patients (patient 2 and patient 3) treated with DMSO or MA-5 in comparison with the OCR (left) and ECAR (right). The data represent the mean ± SD.

**Figure S5**

Effect of MA-5 on mtROS. MA-5 improved ROS production by BSO. DHR123 staining of sIBM myoblasts treated with DMSO (upper left), MA-5 (upper right), BSO + DMSO (lower left), or BSO + MA-5 (lower right). The data represent the mean ± SEM. ###*p*< 0.001 (unpaired two-tailed Student’s *t*-test versus DMSO); \*\**p*< 0.01, \*\*\**p*< 0.001 (unpaired two-tailed Student’s *t*-test versus BSO + DMSO). Scale bars = 200 μm.

**Figure S6**

Electron microscopy analysis of sIBM myoblasts. Structural analysis of cristae in sIBM myoblasts analyzed by electron microscopy. sIBM myoblasts were treated with (a) 0.1% DMSO or (b) 10 μM MA-5 for 24 h. The maximal width and length of the cristae were compared by ImageJ software (n=100). Scale bars = 500 nm.

**Figure S7**

Mfn2 expression levels. Mfn2 was assessed at the transcript level in myoblasts from normal controls and sIBM patients.

**Figure S8**

Cell death induced by BSO treatment. (a) sIBM1 myoblasts. (b) sIBM2 myoblasts. (c) sIBM3 myoblasts. The figure on the left shows the cell viability assay, and the figure on the right shows the level of LDH in the culture medium under oxidative stress conditions induced by BSO treatment. The data represent the mean ± SEM. \**p*< 0.05, \*\**p*< 0.01 and \*\*\**p*< 0.001 (unpaired two-tailed Student’s *t*-test versus DDW; n=4). The black square indicates a significant increase or decrease compared with DDW.

**Figure S9**

Cell protective effect of MA-5 in sIBM myoblasts. (a) Normal myoblasts. (b) sIBM1 myoblasts. (c) sIBM2 myoblasts. (d) sIBM3 myoblasts. The figure on the left shows the cell viability assay, and the figure on the right shows the level of LDH in the culture medium under oxidative stress conditions induced by BSO treatment. The BSO concentration required to reach approximately 50-80% cell death varied with the type of myoblast. The data represent the mean ± SEM. ##*p*< 0.01 and ###*p*< 0.001 (unpaired two-tailed Student’s *t*-test versus control myoblasts; n=4); \**p*< 0.05, \*\**p*< 0.01 and *** *p*< 0.001 (unpaired two-tailed Student’s *t*-test versus BSO+DMSO; n=4). The black square indicates a significant increase or decrease compared with DMSO.

**Figure S10**

Electron microscopy analysis of sIBM fibroblasts. Structural analysis of cristae in sIBM fibroblasts analyzed by electron microscopy. sIBM fibroblasts were treated with (a) 0.1% DMSO or (b) 10 μM MA-5 for 24 h. The maximal width and length of the cristae were compared by ImageJ software (n=100). Scale bars = 200 nm.

